# Immunocompetent murine model of Ewing sarcoma reveals role for TGFβ inhibition to enhance immune infiltrates in Ewing tumors during radiation

**DOI:** 10.1101/2024.05.07.592974

**Authors:** Jessica D. Daley, Elina Mukherjee, A Carolina Tufino, Nathanael Bailey, Shanthi Bhaskar, Nivitha Periyapatna, Ian MacFawn, Sheryl Kunning, Cynthia Hinck, Tullia Bruno, Adam C. Olson, Linda M. McAllister-Lucas, Andrew P. Hinck, Kristine Cooper, Riyue Bao, Anthony R. Cillo, Kelly M. Bailey

**Affiliations:** Department of Pediatrics, University of Pittsburgh School of Medicine, Pittsburgh, PA, USA; Department of Pathology, University of Pittsburgh School of Medicine, Pittsburgh, PA, USA; UPMC Hillman Cancer Center, Pittsburgh, PA, USA; Department of Immunology, University of Pittsburgh School of Medicine, Pittsburgh PA, USA; Department of Structural Biology, University of Pittsburgh School of Medicine, Pittsburgh, PA, USA; Department of Radiation Oncology, University of Pittsburgh School of Medicine, Pittsburgh, PA, USA; UPMC Hillman Cancer Center Biostatistics Facility, Pittsburgh, PA, USA; UPMC Hillman Cancer Center Bioinformatics, Pittsburgh, PA, USA

## Abstract

Ewing sarcoma (ES) is an aggressive cancer diagnosed in adolescents and young adults. The fusion oncoprotein (EWSR1::FLI1) that drives Ewing sarcoma is known to downregulate *TGFBR2* expression (part of the TGFβ receptor). Because *TGFBR2* is downregulated, it was thought that TGFβ likely plays an inconsequential role in Ewing biology. However, the expression of TGFβ in the Ewing tumor immune microenvironment (TIME) and functional impact of TGFβ in the TIME remains largely unknown given the historical lack of immunocompetent preclinical models. Here, we use single-cell RNAseq analysis of human Ewing tumors to show that immune cells, such as NK cells, are the largest source of TGFβ production in human Ewing tumors. We develop a humanized (immunocompetent) mouse model of ES and demonstrate distinct TME signatures and metastatic potential in these models as compared to tumors developed in immunodeficient mice. Using this humanized model, we study the effect of TGFβ inhibition on the Ewing TME during radiation therapy, a treatment that both enhances TGFβ activation and is used to treat aggressive ES. Utilizing a trivalent ligand TGFβ TRAP to inhibit TGFβ, we demonstrate that in combination with radiation, TGFβ inhibition both increases ES immune cell infiltration and decreases lung metastatic burden *in vivo*. The culmination of these data demonstrates the value of humanized models to address immunobiologic preclinical questions in Ewing sarcoma and suggests TGFβ inhibition as a promising intervention during radiation therapy to promote metastatic tumor control.

## Introduction

The tumor microenvironment (TME) is a dynamic, interconnected network of tumor cells and other non-malignant supporting elements such as stromal cells, immune cells, cytokines, etc (1). Tumor treatment, such as chemotherapy and radiation, remodels the TME by promoting tumor cell apoptosis, altering the TME cytokine profile, shifting the composition of tumor immune infiltrates, and driving functional dysregulation of immune cells (2–4). The TME can support and promote tumor cell growth, metastases, and survival and paradoxically, in some cases, tumor treatment enhances the tumor-promoting effects of the TME (5–8). Emerging cancer therapeutics are designed to remodel the TME to reduce immunosuppression and promote tumor control (9, 10).

Recent studies of the TME of Ewing sarcoma, an aggressive fusion oncoprotein (EWSR1::FLI1)-driven primary bone tumor of adolescents and young adults, (11–13) revealed the importance of extra-cellular matrix (ECM) remodeling (14) and hypoxia (15, 16) in promoting Ewing tumor cell survival and metastatic potential. An aspect of the Ewing TME that continues to be poorly understood --and not well represented in *in vivo* studies of Ewing sarcoma--is Ewing immunobiology. This, in part, is due to the historical lack of a syngeneic or transgenic (immunocompetent) mouse models of this cancer. Despite international efforts over decades to develop a transgenic mouse model of Ewing sarcoma (17, 18), attempts to induce expression of EWSR1::FLI1 at the right level, in the right cell, and in the correct developmental state have proved very challenging and ultimately unsuccessful. In 2022, Vasileva et. al developed an immunocompetent zebrafish model of Ewing sarcoma; characterization of the tumor immune microenvironment of this model has not yet been explored (19).

Our current understanding of the Ewing tumor immune microenvironment is largely derived from the analysis of limited human patient samples. Primary, pre-treatment Ewing tumor FFPE (formalin-fixed paraffin-embedded) specimens from the bone demonstrate an overall low immune infiltrate (20). Recent work from our group utilized fresh tumor and paired peripheral blood specimens from patients with Ewing sarcoma to conduct single-cell RNAseq analysis of CD45+ (leukocyte common antigen) cells infiltrating tumors in order to advance the understanding of the evolution of Ewing immunobiology with disease progression. Ewing tumors demonstrated increased immune cell infiltration at the time of relapse in comparison to the time of original diagnosis, and intracellular communication analyses identified CD14+CD16+ macrophages as drivers of immune infiltration. We found that, in contrast to our parallel analyses of osteosarcomas, the mechanism of this effect was less dependent on CXCL10/CXCL12 (21). The specific factors in the tumor microenvironment that modulate Ewing tumor immunobiology, including cytokine expression, remain largely unknown.

In the current study, using single-cell RNAseq analysis, we identify high expression of *TGFB1* specifically in the immune compartment of human Ewing tumors. We develop and utilize a CD34+ humanized mouse model of Ewing sarcoma (human immune cells plus human tumor cells) to study the impact of an intact immune system on Ewing response to radiation therapy, a treatment modality that is commonly used for patients with metastatic and relapsed Ewing sarcoma (22–24). To understand immune contexture-dependent contributions to the Ewing transcriptome, we conducted parallel studies of identical Ewing tumors in immunodeficient NSG mice. We then identify and compare immune-dependent proinflammatory signatures induced by radiation therapy. We successfully target TGFβ, an immunosuppressive cytokine, utilizing a TGFβ TRAP during radiation in these models. Inhibition of TGFβ during radiation in our humanized mouse model of Ewing sarcoma both increases immune infiltration in these classically immune excluded tumors and decreases lung metastases. These data point to the importance of the tumor microenvironment in regulating aspects of Ewing biology and highlights the importance of TGFβ in Ewing sarcoma, despite Ewing tumor cells demonstrating reduced *TGFBR2* (TGF-beta receptor 2) expression (25). This work provides rationale for additional preclinical testing of TGFβ inhibition during radiation therapy for the treatment of aggressive Ewing sarcoma. More broadly, we demonstrate a promising new approach to enhance TME modeling *in vivo* and we establish a platform for additional preclinical testing of agents in Ewing sarcoma that may be impacted by the presence of an intact immune system.

## Materials and Methods

### Cell lines

TC32 (Childhood Cancer Repository, cccells.org) and A673 (ATCC) Ewing cell lines were cultured in glutamine containing, phenol-free RPMI supplemented with 10% FBS. Cell line identification was authenticated by short tandem repeat (STR) profiling at the University of Arizona Genetics Core. Mycoplasma contamination was monitored using the MycoAlert PLUS Mycoplasma Detection Kit (Lonza, catalog number LT07-701). HLA-A2 status of Ewing tumor cell lines was determined.

### Mouse models and establishment of Ewing tumors

Protocols for *in vivo* mouse experiments were approved by the Institutional Animal Care and Use Committee (IACUC) at the University of Pittsburgh under the reference #22010361. Hu-CD34+ mice were purchased from Jackson laboratory (strain number 705557). The hu-CD34+ strain is an NSG mouse that is engrafted with CD34+ cord-blood derived hematopoietic stem cells. Immunodeficient NSG mice were also purchased from Jackson laboratory (strain number 005557). For establishment of tumors in both the Hu-CD34+ mice and the NSG mice, 5 x 10 ^5 TC32 or A673 cells (mixed in a 1:1 solution of growth factor reduced Geltrex [Life Technologies, catalog number A1413202] with PBS) were injected into the mouse in a peritibial location.

### Tumor treatment with radiation therapy

Lab members completed radiation safety training at The University of Pittsburgh. Mice bearing Ewing tumors received a single dose of 4 or 5 Gy (dose determine by prior *in vitro* testing) with an X-RAD 320 (Precision X-ray Inc, N. Branford, CT USA) using Filter 2 (1.5mm Al + 0.25mm Cu +0.75mm Sn). A full body shield with window to expose the lower extremity (Precision X-ray, part number XD1907-2021) was used to isolate the peri-tibial tumors for focal radiation therapy and minimize off-tumor effects.

### Mouse treatment with RER and plasma testing of TGFβ inhibition

Endotoxin-free RER was prepared as previously described (37) and was delivered at a dose of 50 micrograms per day via IP injection, as previously described (50). Mice receiving vehicle control underwent IP injection with PBS (50 microL daily). Treatment with either RER or PBS started on day 1 of radiation and continued for 6 total doses with sacrifice occurring on day 7. Plasma isolated from mice at time of sacrifice was analyzed using ELISA (Abcam Inc, catalog number ab1006447) for quantification of TGFβ1.

### FFPE preparation and immunohistochemistry

At time of sacrifice, tumors and lungs were collected. For tissue histology, a portion of the tumor as well as both lungs were placed in formalin for at least 24 hrs prior to paraffin embedding. The University of Pittsburgh PBC (Pitt Biospecimen Core) acquired 5 micron serial sections (interval of 500 microns) of the blocks and performed H&E staining. The UPMC Department of Pathology clinical IHC protocol was used for CD99 and NKX2.2 staining (markers of Ewing sarcoma).

### Examination of lung metastases

Serial lung sections (5 slides per mouse) were reviewed an independent pathologist blinded to the treatment conditions. Independent metastatic lesions on each slide was counted as a single metastatic focus.

### Blood and tumor processing and flow cytometry analysis

Blood was collected from mice at time of sacrifice. Whole blood was spun at 400xG for 10 minutes with brake off. Plasma was collected and immediately stored at −80C for future analysis. Red cell lysis was performed on the remaining cell pellet using BD Pharmlyse buffer (BD Biosciences, catalog number 555899) using the manufacturer’s instructions. The remaining cell pellet was washed and resuspended in PBS. Mouse tumors were manually dissected into 1 mm^2^ pieces with a scalpel and incubated for 15 minutes at 37°C in 5% CO2 in 5mL of serum free, phenol free RPMI and 50 mg/mL Liberase DL (Millipore Sigma, Catalog No. 5466202001). Single-cell suspensions were then passed over a 70 μmol/L filter to remove debris. Both PBMCs and tumor cell suspensions were stained with conjugated antibodies for 20 minutes in the dark. The following antibodies were used for surface staining: Human CD163 BV650 (BD Biosciences 563888), Human CD25 BV711 (Biolegend Inc 302636), Human CD20 BV750 (BD Biosciences 747062), Human CD8 BUV496 (BD Biosciences 612942), Human CD45 BUV395 (BD Biosciences 563792), Human CD14 BUV737 (BD Biosciences 612763), Human CD4 BUV563 (BD Biosciences 741353), Human CD103 BUV615 (BD Biosciences 751285), Human CD3 Spark Blue 550 (Biolegend Inc 344852), Human CD11b PerCP/Cyanine5.5 (Biolegend Inc 301328), CD56, Human CD19 APC/Firetrade 810 (Biolegend Inc 302272), Human HLA-DR Alexa Fluor 488 (Biolegend Inc 307620), Human CD16 PE/Dazzletrade 594 (Biolegend Inc 302054), Human CD33 PE (Biolegend Inc 366608), Human Ki-67 BV421 (Biolegend Inc 350506), and Human FoxP3 EF450 (Invitrogen 48477642). Cells were stained with viability dye Zombie NIRtrade Fixable Viability Kit (Biolegend Inc, catalog number 423105). Stained cells were fixed using eBioscience Fixation/Permeabilization Concentration (Invitrogen, catalog number 00-5123-43). The data were acquired on the Cytek Aurora spectral flow cytometer and analyzed using the Flow Jo software v10.9.0. Additional analysis was performed utilizing Cytobank Software (beckman.com).

### Tumor RNA extraction and bulk RNA-sequencing

A portion of tumors isolated from mice at sacrifice were flash frozen using liquid nitrogen. RNA isolation was performed on this tumor section using Qiagen QIAshredder (Qiagen, cat no: 79656) and Qiagen RNEasy Plus Micro Isolation Kit. (Qiagen, cat no: 74034). RNA concentration was measured using Nanodrop (Thermo Scientific). mRNA library preparation and sequencing were performed at the Health Sciences Sequencing Core at Children’s Hospital of Pittsburgh with a 2×50 paired end read length and 40M reads per sample.

### RNA Sequencing Library Generation

RNA was assessed for quality using an Agilent TapeStation 4150/Fragment Analyzer 5300 and RNA concentration was quantified on a Qubit FLEX fluorometer. Libraries were generated with the Illumina Stranded mRNA Library Prep kit (Illumina: 20040534) according to the manufacturer’s instructions. Briefly, 1000 ng of input RNA was used for each sample. Following adapter ligation, 10 cycles of indexing PCR were completed, using IDT for Illumina RNA UD Indexes (Illumina: 20040553-6). Library quantification and assessment was done using a Qubit FLEX fluorometer and an Agilent TapeStation 4150/Fragment Analyzer 5300. Libraries were normalized and pooled to 2 nM by calculating the concentration based off the fragment size (base pairs) and the concentration (ng/μl) of the libraries.

### Library Sequencing

Sequencing was performed on an Illumina NextSeq 2000, using a P3 100 flow cell. The pooled library was loaded at 750 pM and sequencing was carried out with read lengths of 2×58 bp, with a target of 40 million reads per sample. Sequencing data was demultiplexed by the on-board Illumina DRAGEN FASTQ Generation software (v3.10.12).

### Bulk RNAseq gene expression quantification

After quality control, the paired-end reads were pseudoaligned to human reference transcriptome (GRCh38) with Gencode annotation(v28) and summarized at transcript level using kallisto (v0.48.0). Transcript abundance was summarized into gene level using tximport (v1.22.0), Trimmed Mean of M-values (TMM) normalized, and log2-transformed. Genes with low expression (counts per million of mapped reads [CPM] < 3) were removed prior to normalization and statistical comparison.

### Human Ewing tumor processing for Single-cell RNAseq

Viably preserved Ewing tumor biopsy specimens were rapidly thawed, processed into single cell suspensions, and immediately loaded into the 10X Genomics Controller as per the manufacturer’s protocol (as previously described (21)) in order to analyze gene expression signatures of individual cells within the tumor microenvironment (IRB approved STUDY19030108). 10x Genomics library preparation was conducted at The University of Pittsburgh Single Cell Core.

### Processing of single-cell RNAseq data

Following sequencing, raw sequencing files were demultiplex into FASTQ files using bcl2fastq (v.2.20.0) and were aligned to the reference genome GRCh38 using CellRanger (v6.0.1). Following alignment, feature/barcode matrices were read into Seurat (v4.3.0.1) in R (4.2.0). Previously described data from Ewing sarcoma patients (21) was also read into Seurat and a unified analysis object was created. Cells with fewer than 200 genes per cell or with >10% of reads aligning to mitochondrial genes were excluded as part of a quality control workflow.

### Dimensionality reduction and clustering

Dimensionality reduction was performed in Seurat as previously described (21) with the addition of an integration workflow. Briefly, the top 2000 highly variable genes were identified in each sample and SCTransform was used to normalize expression values in each sample individually. Next, anchors were identified, and data integration was performed to reduce technical variation between samples. Following integration, principal component analysis was performed using the 2000 integrated features. Consecutive principal components were heuristically selected based on the variance explained per principal component and the selected principal components were used for the generation of UMAPs and as input for Leiden-based clustering.

### Identification of cell types

Cell types were identified as previously described (21). Briefly, cell types were identified based on the expression of canonical lineage markers including for leukocytes (lymphocytes including T cell subsets, B cells, and NK cells; myeloid cells including monocytes, macrophages, and dendritic cell) tumor populations, osteoclasts, and fibroblasts using markers *PTPRC, CD3D, CD4, CD8A, FOXP3, CD14, FCGR3A, MS4SA1, CD1C, IL3RA, MMP9*, and *COL1A1*. Lymphocyte subsets were further refined by performing dimensionality reduction and clustering on only lymphocyte subsets.

### Differential gene expression detection and pathway enrichment analysis

Genes differentially expressed (DEGs) between groups of interest was identified using Linear Models for Microarray and RNA-Seq Data (limma) voom algorithm with precision weights (v3.50.3), and filtered by FDR-adjusted P<0.05 and fold change ≥ 1.5 or ≤ − 1.5. Pathways enriched in significant DEGs were identified using Enrichr with BioPlanet2019 (accessed August 2023).

### Causal network analysis

Upstream transcriptional regulators and change of direction (activation or inhibition) was predicted from target molecules (encoded by significant DEGs) using Ingenuity Pathway Analysis (IPA ®) (QIAGEN Inc., Germany) with the curated Ingenuity Knowledge Base (accessed August 2023), and filtered at overlap P<0.05 (measuring the enrichment of target molecules in the dataset) and z-score ≥ 1.95 (measuring the predicted activation level of the pathways).

### Digital immune cell population deconvolution

Single-cell RNAseq count matrix and cell type annotations previously published by our group (21) were used to construct gene signature matrix using cibersortx with 1000 permutations and default parameters. The signature was subsequently used for estimating the fraction of immune cell populations in bulk RNAseq data.

### Statistical analysis – RNAseq data

Hypergeometric test was used for pathway enrichment detection. Non-parametric Wilcoxon test was used to compare cell population fractions between groups. BH-FDR was used for adjustment of multiple comparisons. All tests are two-sided unless otherwise noted. FDR was controlled at 0.05 level.

### Statistical analysis

To determine differences in total human CD45+ cell counts a linear model was fit with a fixed effect for treatment group and a random effect for biological replicate. Pairwise differences compared to control only were evaluated using degrees of freedom based on Satterthwaites method. To determine difference in serum levels of TGFB1, paired t-test was performed utilizing Graphpad Prism v.10. To determine the difference in lung metastatic burden, the correlation of counts across slides was adjusted by calculating the average number of lung mets across all five slides for each mouse. A poisson model was fit to the data to model the average count of lung mets across treatment groups. A random effect was included to control for correlation of mice within human donor replicate. Statistical analysis was performed using RStudio (v 2023.09.1).

### Software for schematics

Schematics were created using biorender.com.

## Results

### *TGFB1* expression in human Ewing tumors is predominantly in the immune cell compartment

TGFβ is an immunosuppressive cytokine that can prevent effective anti-tumor immune responses and remodel the extra-cellular matrix toward a pro-tumor phenotype (13, 14). Increased TGFβ pathway signaling is noted in Ewing tumors that demonstrate a pro-metastatic, aggressive phenotype (26). Recent work also demonstrates that TGFβ is elevated in the serum of patients with Ewing sarcoma, as compared to healthy controls (27). Strategies to negate the pro-tumorigenic impact of TGFβ and to enhance the efficacy of cellular therapies are currently being tested in clinical trials for relapsed Ewing sarcoma (NCT05634369) (28).

Currently, little is understood regarding the contribution of individual cell types in the TME to TGFβ expression. Thus, we sought to analyze TGFβ expression in human Ewing tumors on a single cell level. We conducted single-cell RNA sequencing (scRNAseq) on fresh (21) and viably preserved human Ewing tumors acquired through an IRB-approved protocol (Fig 1a). Resulting cell clusters were identified based on transcriptomic profiles (Fig 1b). We then queried the expression of *TGFB1* in these cell populations and find that *TGFB1* is most highly expressed by immune cell populations in human Ewing tumors (Fig 1c), with low expression from tumor and stromal cell populations. Specifically, expression of *TGFB1* is highest in natural killer cells, CD4+ T regulatory cells, CD8+ T cells, and CD14+CD16-monocytes isolated from human Ewing tumors (Fig 1d). These findings confirm that TGFβ is expressed in the Ewing tumor microenvironment and almost exclusively in the immune cell compartment. This data also suggest that in order rigorously determine the effects of TGFβ on the Ewing tumor microenvironment, an immunocompetent preclinical model is necessary.

**Figure 1.**
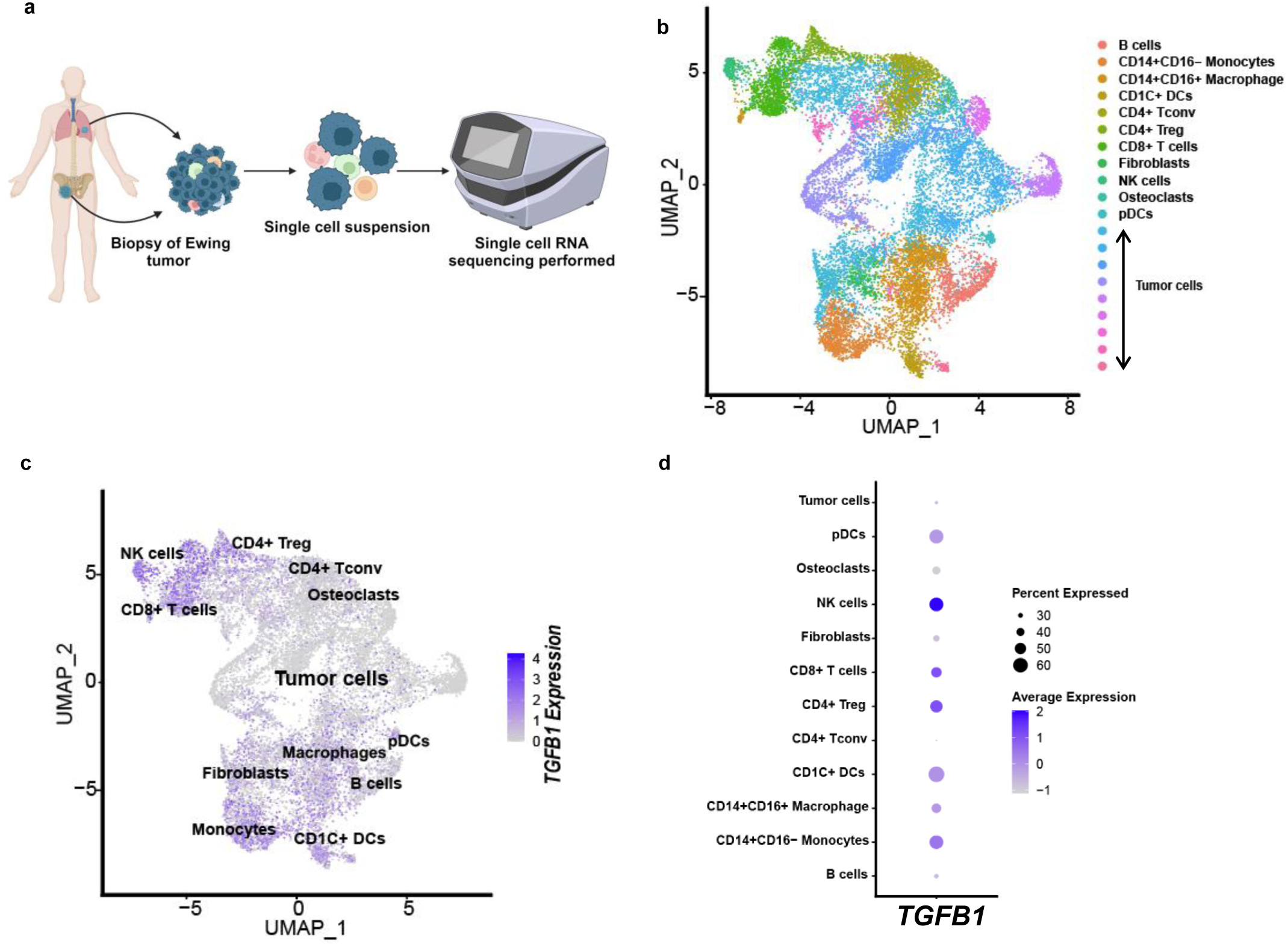
*TGFB1* is predominantly expressed by immune cell populations in Ewing tumors. **a** Experimental overview. **b** UMAP embedding of 17,610 cells from tumors of Ewing sarcoma patients. **c** Expression of *TGFB1* across immune, non-immune, and tumor populations from the six patient samples shown in **b**. Log2 expression levels of *TGFB1* are visualized across all cells (left panel). Inferred cell types from all cells and patients are shown (right panel). *TGFB1* is expressed across numerous lymphocyte (CD4+ Treg, CD8+ T cell, and NK cells) and myeloid cell (monocytes, macrophages and DCs) populations in the Ewing tumor microenvironment. **d** Dot plot showing the mean *TGFB1* expression levels (colors of the dot) and the frequency of cells that expression *TGFB1* (size of the dot) across all cells from the tumor microenvironment.

### Ewing tumors developed in a hu-CD34+ model are biologically and transcriptionally distinct from tumors developed in an immunodeficient model

As a next step toward elucidating the role of TGFβ in the Ewing tumor immune microenvironment, we developed and validated an immunocompetent model for *in vivo* studies of Ewing tumors. To develop this model, we utilized hu-CD34+ humanized mice. The hu-CD34+ murine model is derived from an immunocompromised, NSG mouse that undergoes transplantation utilizing human CD34+ cord blood derived stem cells (29, 30) resulting in reconstitution with multiple CD45+ human immune cell lineages including B cells, T cells, and myeloid cells (Supplemental Fig 1). Ewing tumors were established in an orthotopic peri-tibial location (Fig 2a). Resulting tumors were confirmed to be Ewing sarcoma with positive immunohistochemistry staining for CD99, a membrane protein expressed uniformly in Ewing tumors, and nuclear staining of NK homeobox 2 (NKX2.2), a transcription factor upregulated by the EWSR1::FLI1 (Fig 2b). Of note, spontaneous pulmonary metastases develop in this model (Fig 2c), a finding which recapitulates the situation in humans, as the lung is the most common site of metastasis in patients with Ewing sarcoma (31). Peripheral blood was also collected from mice at sacrifice and we demonstrate that the human immune compartment persists in the peripheral blood of these mice throughout the duration of the experiment (Fig 2d). Immunophenotyping was also performed on single cell suspensions of Ewing tumors established in the hu-CD34+ model and demonstrates low baseline infiltration of hu-CD45+ cells in Ewing tumors (Fig 2e), a result that is similar to the level of immune infiltration of Ewing bone tumors noted in patients at baseline (32, 33).

**Figure 2.**
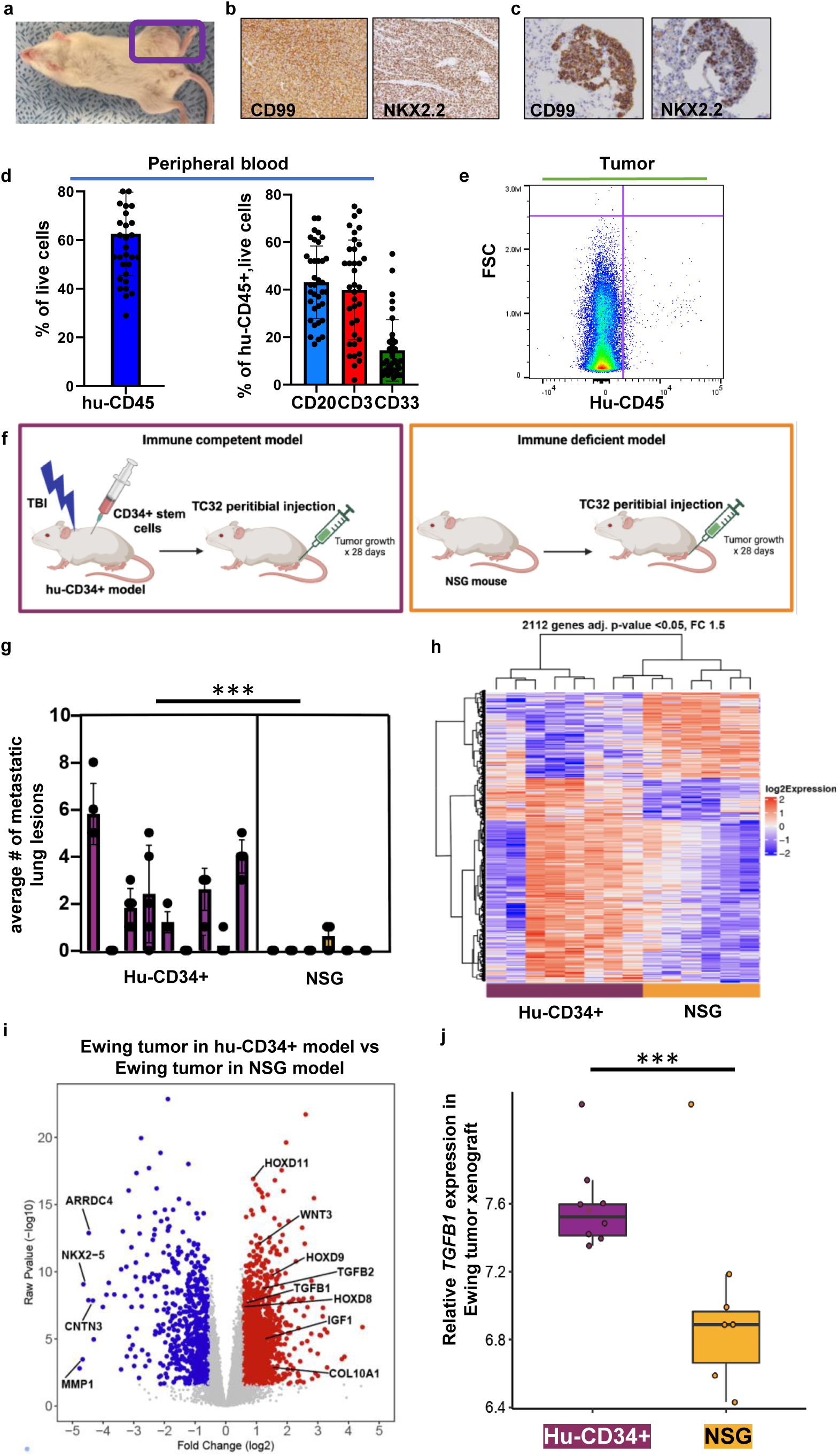
Ewing tumors developed in hu-CD34+ model are biologically and transcriptionally distinct from tumors developed in an immunodeficient model. **a** Orthotopic Ewing tumors are successfully established in a humanized (hu-CD34+) mouse. **b** Representative CD99 and NKX2.2 IHC staining of Ewing tumors developed in a hu-CD34+ model, magnification 200x. **c** Representative CD99 and NKX2.2 IHC staining of lung metastases that develop in the hu-CD34+ Ewing model, magnification 200x. **d** Immuno-phenotyping performed on peripheral blood mononuclear cells obtained from mice bearing TC32 Ewing tumors at sacrifice following tumor growth demonstrates persistence of engrafted human immune cells. HuCD45 represents % of all peripheral lymphocytes that are huCD45 positive. The percent of hu-CD45 cells that express CD20, CD3, and CD33 (representing B cells, T cells and myeloid cells respectively) are shown. n = 36 mice. Error bars represent standard deviation. **e** Representative flow cytometry plot demonstrates low (1.24% of all cells in the tumor) baseline huCD45+ infiltration in Ewing tumors developed in hu-CD34+ mice. **f** Schematic depicting hu-CD34+ versus NSG mouse models used in these experiments. **g** Lungs were harvested from hu-CD34+ or NSG mice harboring TC32 Ewing cell tumors, and the presence of metastatic foci per section was determined by an independent pathologist. Each bar represents average number of metastatic lesions per slide in individual mouse. *** = p< 0.05. Error bars represent standard deviation. **h** Bulk RNA sequencing was performed on TC32 Ewing tumors developed in either hu-CD34+ versus NSG mouse models. A heat map depicting differential gene expression of tumors established in hu-CD34+ mice vs NSG mice is shown. **i** Volcano plot of differentially expressed genes in TC32 tumors established in hu-CD34+ mice vs NSG mice. **j** Relative *TGFB1* expression in Ewing tumors developed in NSG mice and hu-CD34+ mice was determined, ***=p<0.05. Error bars represent standard deviation.

Having demonstrated the feasibility of utilizing a hu-CD34+ Ewing model, we next sought to understand the transcriptomic and biologic differences noted in Ewing tumors established in hu-CD34+ versus NSG (immunodeficient) mice. Tumors were established in both the hu-CD34+ immunocompetent mouse model and an NSG immunodeficient murine model utilizing the human TC32 Ewing sarcoma cell line (Fig 2f). Tumors were allowed to develop for approximately four weeks prior to sacrifice, at which time both tumors and lungs were collected. Lung tissue was formalin-fixed and paraffin-embedded and the resulting blocks underwent serial sectioning. Serial lung sections were reviewed independently by a pathologist to determine the burden of lung metastases. We find significantly more lung metastases present in the hu-CD34+ mice as compared to NSG mice (Fig 2g). To examine transcriptional differences between Ewing tumors developed in hu-CD34+ versus NSG mice, bulk RNA sequencing was performed on RNA isolated from flash frozen tumors. Unsupervised clustering of the transcriptional profiles across all samples indicate that Ewing tumors developed in the hu-CD34+ and immunodeficient NSG models cluster separately (Fig 2h). A total of 2112 differentially expressed genes (fold change >1.5, p value <0.05) were identified in tumors developed in a humanized model versus NSG model (Fig 2i). Upregulated gene transcripts in Ewing tumors developed in the hu-CD34+ models compared to the NSG model include *Wnt3,* Hox gene family members, and collagen family genes, some of which have been previously reported to be pro-tumorigenic in Ewing sarcoma (14). Tumors grown in hu-CD34+ mice demonstrate upregulation of pathways driving collagen biosynthesis and extracellular matrix (ECM) remodeling (Supp. Fig 2), when compared to tumors from immunodeficient mice.

In addition, we hypothesized that *TGFB1* expression would be increased in tumors developed in an immunocompetent model since our human Ewing tumor data demonstrated that *TGFB1* is expressed predominantly in the immune compartment (Fig 1). We compared the relative expression of *TGFB1* in Ewing tumors derived developed in the hu-CD34+ versus NSG model and indeed find that *TGFB1* is significantly upregulated in the Ewing tumors developed in the hu-CD34+ model (Fig 2j). Taken together, these data demonstrate that Ewing tumors developed in an immunocompetent, humanized murine model have increased expression of important modulators of the tumor microenvironment, including *TGFB1*, and increased metastatic potential, as compared to tumors developed in NSG, immunodeficient models.

### Post-radiation therapy transcriptional remodeling in Ewing tumors is immune-contexture dependent

Previous investigations in epithelial-derived cancers (non-sarcomas) have demonstrated an important role for TGFβ in modulating the net response to cancer therapies (34, 35). Specifically, TGFβ is known to be activated in the TME following radiation therapy, and when present, can blunt the anti-tumor immune response. Importantly, radiation therapy is an essential local control modality for patients with unresectable, metastatic, or relapsed Ewing sarcoma. Recent clinical trials have demonstrated that radiation therapy for localized Ewing sarcoma is an equivocally effective local control method for Ewing tumors that cannot be surgically resected (36). However, the effect of radiation on the tumor immune microenvironment of Ewing sarcoma is not well understood.

We sought to determine the immunomodulatory effect of radiation therapy on the Ewing tumor microenvironment. To do this, Ewing tumors established using TC32 Ewing sarcoma cells in both our hu-CD34+ mouse model and in an NSG model were allowed to develop for 3 weeks. Tumors were then treated with 5Gy radiation therapy as a single fraction (versus 0Gy control) and tumors were harvested 7 days from the start of treatment. A second human Ewing sarcoma cell line, A673 was also tested (Fig 3a). RNA isolated from flash frozen tumors was subjected to bulk RNA-seq and data were analyzed. Transcriptional data underwent unsupervised clustering and demonstrated that transcriptional profiles of Ewing tumors treated with radiation and developed in the hu-CD34+ model cluster separately from Ewing tumors treated with radiation and developed in immunodeficient mice (Supp. Fig. 3). When comparing the transcriptome of tumors post radiation vs untreated tumors, we see more robust transcriptional modulation occurs after radiation in tumors developed in the hu-CD34+ mice as compared to tumors developed in NSG mice (Fig 3b). A total of 2281 significantly differentially expressed genes (p value <0.05, fold change >= 1.5) were identified post-radiation (as compared to no radiation) in hu-CD34+ tumors utilizing the TC32 cell line. Specifically, we find upregulation of genes associated with antigen presentation (*HLA-A, HLA-B, HLA-C*) following tumor treatment with radiation therapy. In contrast, in the NSG model we note fewer transcriptional changes following radiation, with a total of 296 significantly differentially expressed genes (p value <0.05, fold change >= 1.5). We also observe this effect of an intact immune system on radiation-induced transcriptional changes when examining tumors established from the A673 cell line as well (Supp. Fig. 4). Pathway analysis was also performed on RNAseq data from both TC32 tumors and A673 developed in the hu-CD34+ model and significant (p<0.05) upregulation of pathways involved in interferon signaling, antigen presentation, and T cell receptor-induced apoptosis were noted (Supp Fig 5). These findings clearly demonstrate the importance of an intact immune system when preclinically studying the impact of radiation therapy on the Ewing sarcoma TME.

**Figure 3.**
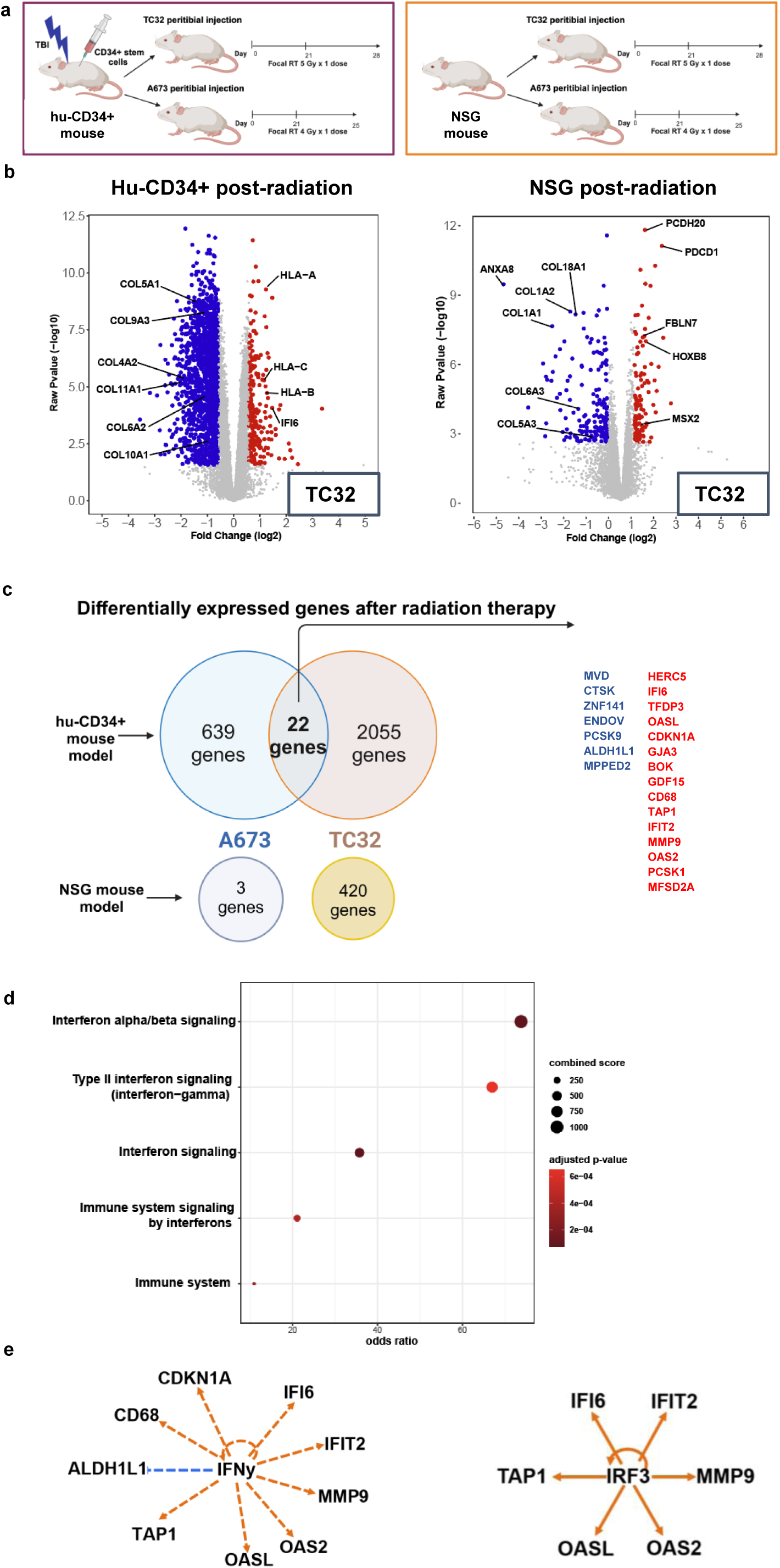
Radiation therapy modulates the Ewing tumor transcriptional profile in an immune contexture-dependent manner. **a** Experimental overview of Ewing tumors established in hu-CD34+ and NSG mouse models followed by treatment with radiation therapy. **b** RNA was extracted from tumors at the experiment termination and bulk RNA sequencing was performed. Volcano plots demonstrating differential gene expression after radiation therapy of TC32 Ewing tumors developed in hu-CD34+ or NSG mice. **c** Venn diagram depicting the number of significantly differentially expressed genes post-radiation therapy when Ewing tumors are developed in a hu-cD34+ versus NSG model. **d** Pathway analysis performed on the shared DEGs after radiation therapy in immunocompetent mouse model utilizing EnrichR bioplanet 2019 library. **e** Upstream analysis utilizing Ingenuity Pathway Analyzer performed on shared gene lists identify IFNy and IRF3 as predicted upstream regulators.

To further evaluate the contribution of an immunocompetent model on radiation-induced transcriptional changes in Ewing tumors, we identified 22 overlapping genes that were differentially expressed following radiation therapy in tumors developed from both TC32 and A673 cell lines in hu-CD34+ mice. No overlapping differentially expressed genes after radiation therapy were identified in tumors developed in immunodeficient mice (Fig 3c). Pathway analysis of this overlapping geneset again identifies upregulation of pro-inflammatory pathways, such as interferon signaling (Fig 3d). We then assessed predicted upstream regulators of these findings utilizing Ingenuity Pathway Analysis. Both IFNγ and IRF7 were predicted to be upstream regulators of the overlapping gene signatures induced by radiation therapy (Fig 3e). Together, these data demonstrate that pro-inflammatory pathways, including interferon signaling, are upregulated following radiation in Ewing sarcoma. When considering the hu-CD34+ *in vivo* data in the context of the human Ewing tumor data presented in Figure 1 (demonstrating an abundance of immunosuppressive TGFβ in the Ewing TME), we next sough to determine the impact of TGFβ inhibition on the Ewing TME following radiation therapy.

### The trivalent ligand TRAP (RER) effectively depletes TGFβ *in vivo*

In order to investigate the role of TGFβ in modulating the Ewing tumor immune microenvironment following radiation, we tested the ability of RER, a trivalent ligand TRAP for TGFβ, to inhibit TGFβ in our hu-CD34+ Ewing model. RER consists of the endoglin-like domain of the rat TGFβ co-receptor betaglycan (BGE, or E) sandwiched between the human TGFβ type II receptor extracellular domains (RII, or R) on both the N- and C terminus, as previously described (37). As illustrated, RER binds and inhibits TGFβ dimers, disabling their ability to serve as receptor ligands (Fig 4a). Importantly, the safety and feasibility of TGFβ ligand TRAPs, similar to RER, were recently validated in human trials (38). To assess the efficacy of RER in our model, Ewing tumors were established in hu-CD34+ mice utilizing the TC32 cell line. Mice were treated with either RER (50 mcg) or vehicle control (PBS) daily for 1 week prior to mouse sacrifice and serum collection. Analysis of mouse serum by TGFβ1 ELISA (enzyme-linked immunosorbent assay) demonstrates that RER effectively depletes systemic TGFβ1 in the hu-CD34+ mouse model (Fig 4b). Next, to determine if RER treatment alters TGFβ signaling in the Ewing tumor microenvironment, we examined the transcriptional profiles of Ewing tumors treated with radiation and RER or vehicle control. Ingenuity Pathway Analysis of bulk RNAseq data identified *TGFB1* to be predicted to be significantly inhibited upstream from the differentially expressed genes identified. These data demonstrate that RER is a successful, potent inhibitor of TGFβ in our *in vivo* model, allowing for both systemic as well an intra-tumoral TGFβ inhibition.

**Figure 4.**
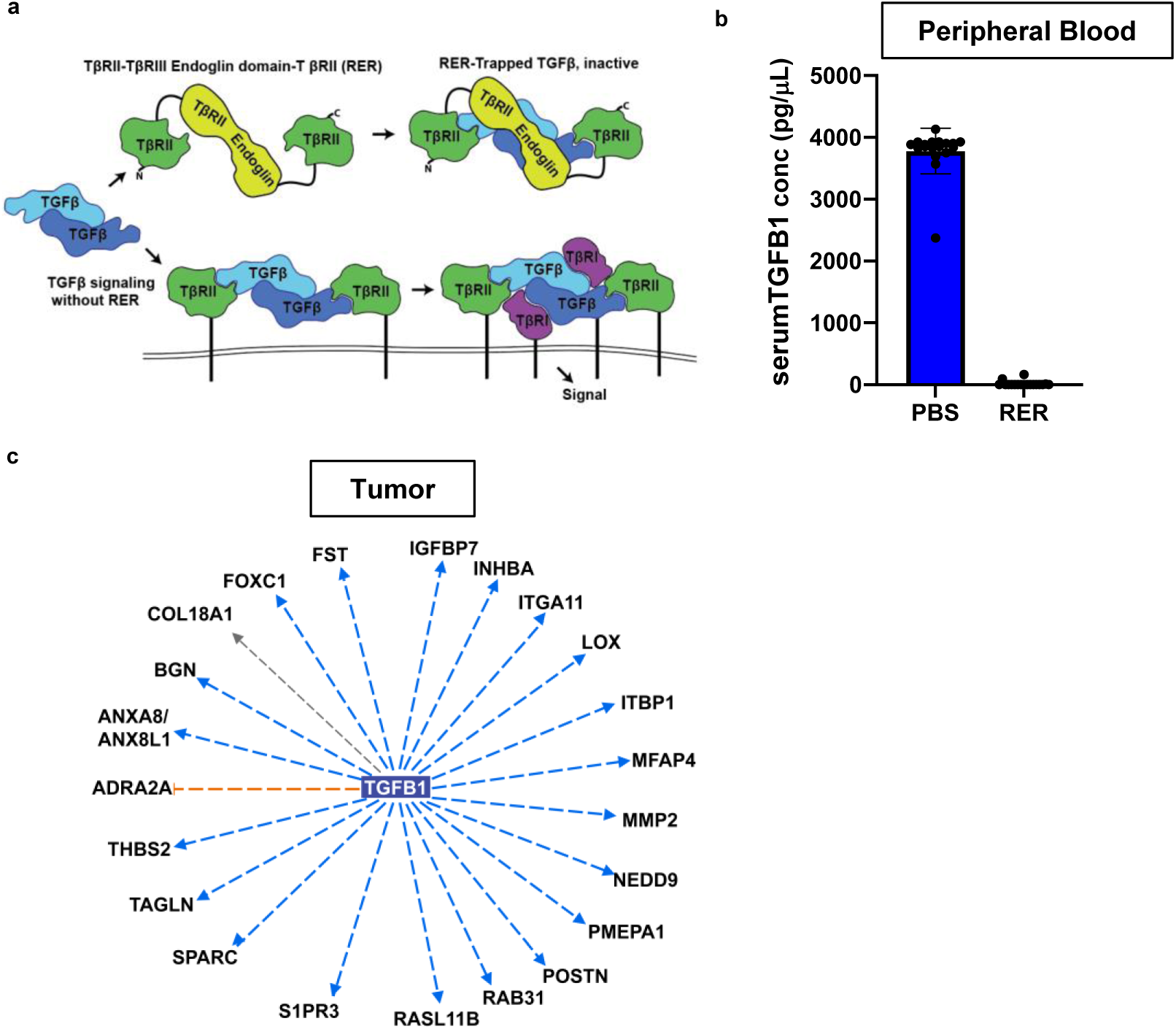
The trivalent ligand TRAP (RER) effectively depletes TGFβ *in vivo.* **a** Schematic of the mechanism of action of TGFβ1 ligand TRAP RER and interaction with TGBRII receptor. **b** Comparison of TGFβ1 levels (determined by ELISA) in the serum of mice at time of sacrifice after receiving IP injections of RER or PBS vehicle control. *** = P<0.05. Error bars represent standard deviation. **c** Bulk RNA sequencing was performed on tumors developed in hu-CD34+ mice that underwent treatment with radiation therapy on day 1 and received either RER or PBS on days 1-7. Ingenuity pathway analysis performed on differentially expressed genes (DEGs) in tumors from mice receiving RER during radiation therapy demonstrates *TGFB1* is predicted to be inhibited upstream of the DEGs. Genes in the outer circle are differentially expressed, blue lines indicate that the genes whose downregulation predict *TGFB1* inhibition.

### TGFβ inhibition during radiation therapy increases immune infiltration of Ewing tumors

After identifying an effective method to inhibit TGFβ *in vivo* (RER), we next focused on understanding the functional impact of TGFβ inhibition during radiation on Ewing tumors in our hu-CD34+ model. We first investigated the effect of TGFβ inhibition during radiation therapy on immune infiltration into Ewing tumors. Ewing tumors were established in the hu-CD34+ model and mice were treated in one of four groups: 1) vehicle control, no radiation, 2) vehicle control and radiation, 3) RER, no radiation, and 4) RER and radiation (Fig 5a). At the time of sacrifice, tumors were harvested, processed into single cell suspensions, and immunophenotyped by flow cytometry. Analysis of the human CD45+ leukocytes infiltrating the Ewing tumors revealed that tumors from mice treated with RER in combination with radiation therapy demonstrate a significant (p<0.05) increase in tumor infiltrating leukocytes as compared to untreated Ewing tumors (Fig 5b). Analysis of immune cell sub-populations demonstrates no significant difference in the ratio of CD3+ T cell and CD3-(non-T cell) populations between each experimental group (Fig 5c). In the CD3+ T cell population we also identify no significant difference in CD4+ or CD8+ T cell populations between treatment groups (Fig 5d). We further determine that in all tumors analyzed, the predominant immune cell population is CD3-, with a predominant myeloid population that is CD14+ and CD14+CD11c+ (Fig 5e, Supp. Fig 6). The CD14+CD11c+ immunophenotype is a signature of myeloid derived suppressor cells (39). Our finding that TGFβ inhibition during radiation significantly increases the total number of hu-CD34+ cells infiltrating Ewing tumors suggests that the immune microenvironment of Ewing tumors is modifiable and that this intervention could be utilized as a tool to enhance total immune cell infiltration in human Ewing tumors.

**Figure 5.**
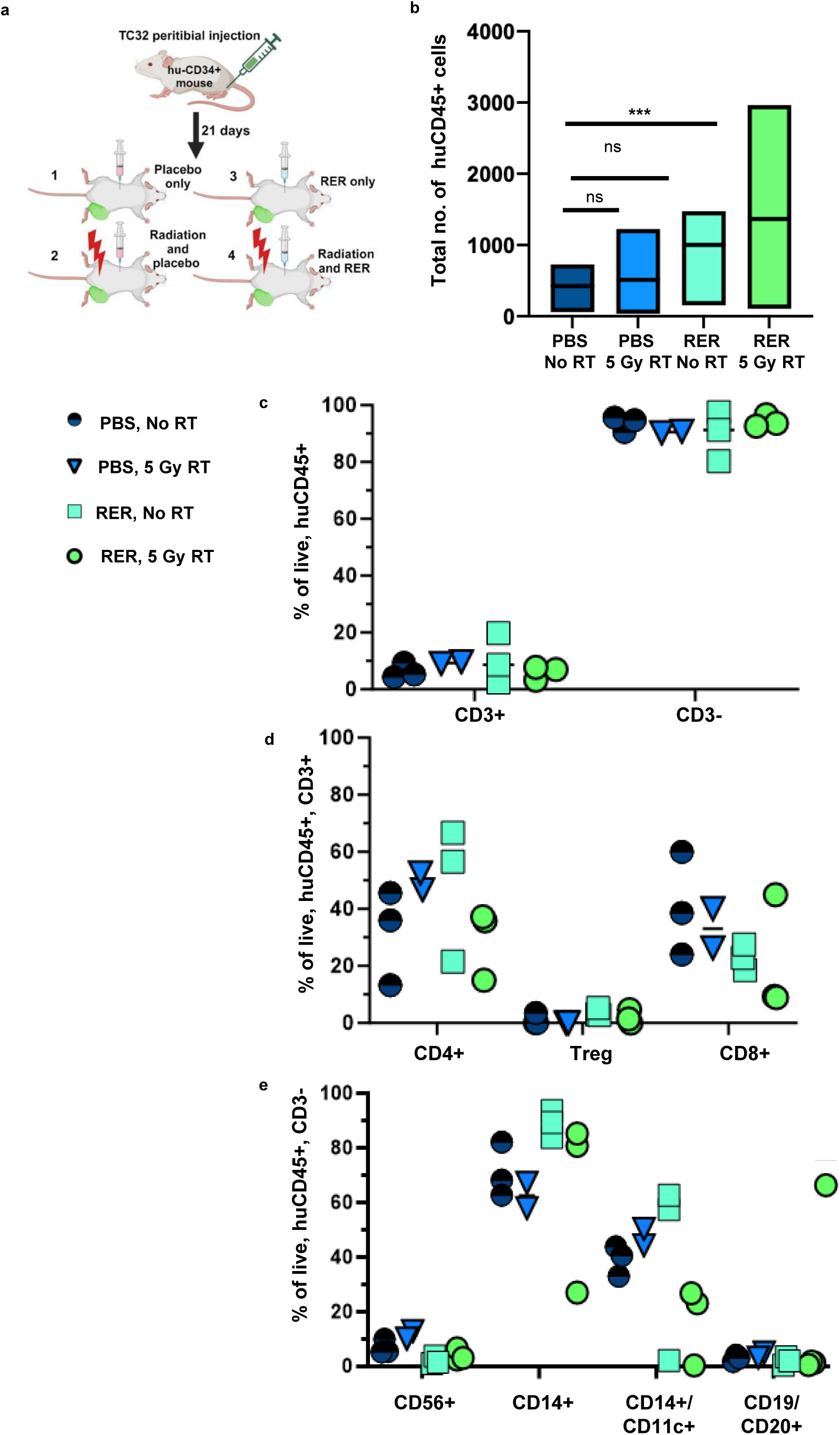
TGFβ inhibition during radiation therapy increases immune infiltration of Ewing tumors. **a** Experimental schematic depicting the four treatment groups. **b** Immunophenotyping was performed on single cell suspensions derived from TC32 Ewing tumors established in hu-CD34+ mice. The total number of huCD45+ cells was determined in each tumor. Box plot represents average total hu-CD45+ count across three biologic replicates, lines represent the mean, and upper and lower limits of the box represents the upper and lower quartiles respectively. *** denotes p<0.05, ns = p>0.05. **c** Immunophenotyping was performed on huCD34+ tumor single cell suspensions. Samples were stained using a multispectral panel (see methods). Samples were analyzed on a Cytek Aurora spectral flow cytometer and unmixed. Files (.fcs) were imported into FlowJo for manual gating. The manual gating strategy is available in Supplemental Figure 6.

### TGFβ inhibition during radiation therapy suppresses the Ewing sarcoma metastatic phenotype

Beyond immunosuppression, TGFβ is also reported to promote tumor metastases(26). It has been shown that subpopulations of Ewing cells termed “EWS::FLI1 low” represent an aggressive population of Ewing cells that demonstrate increased TGFβ signaling. These ‘EWS-FLI1 low” cells regain expression of TGFBR2, one of the two essential TGFβ signaling receptors that is normally repressed by EWSR1::FLI1 (40, 41). Cytokines prevalent in the bone microenvironment, such as Wnt, have been reported to promote an ‘EWS-FLI1 low’ state and enhance Ewing cell responses to TGFβ (42). Having determined that TGFβ inhibition during radiation increases immune cell infiltration into Ewing tumors, we next sought to determine the effect of TGFβ inhibition during radiation on Ewing tumor metastatic potential. To accomplish this, we examined lung tissue from the mice utilized in the experiment shown in Fig 5 (+/− radiation and +/− RER) (Fig 6a). Lungs were isolated at the time of sacrifice and tissue was formalin fixed and paraffin embedded (FFPE). The resulting FFPE blocks underwent serial sectioning and lung tissue underwent review for the presence of metastases by an independent pathologist blinded to treatment conditions. Quantification of these data revealed a significant increase in pulmonary metastases in our humanized model when tumors are treated with radiation alone (Fig 6b) (Supp. Fig 7). In contrast, when mice are treated with RER during radiation, this effect of radiation (an increase in metastasis) is significantly (p<0.05) abrogated, suggesting a role for TGFβ in the development of Ewing tumor lung metastases.

**Figure 6.**
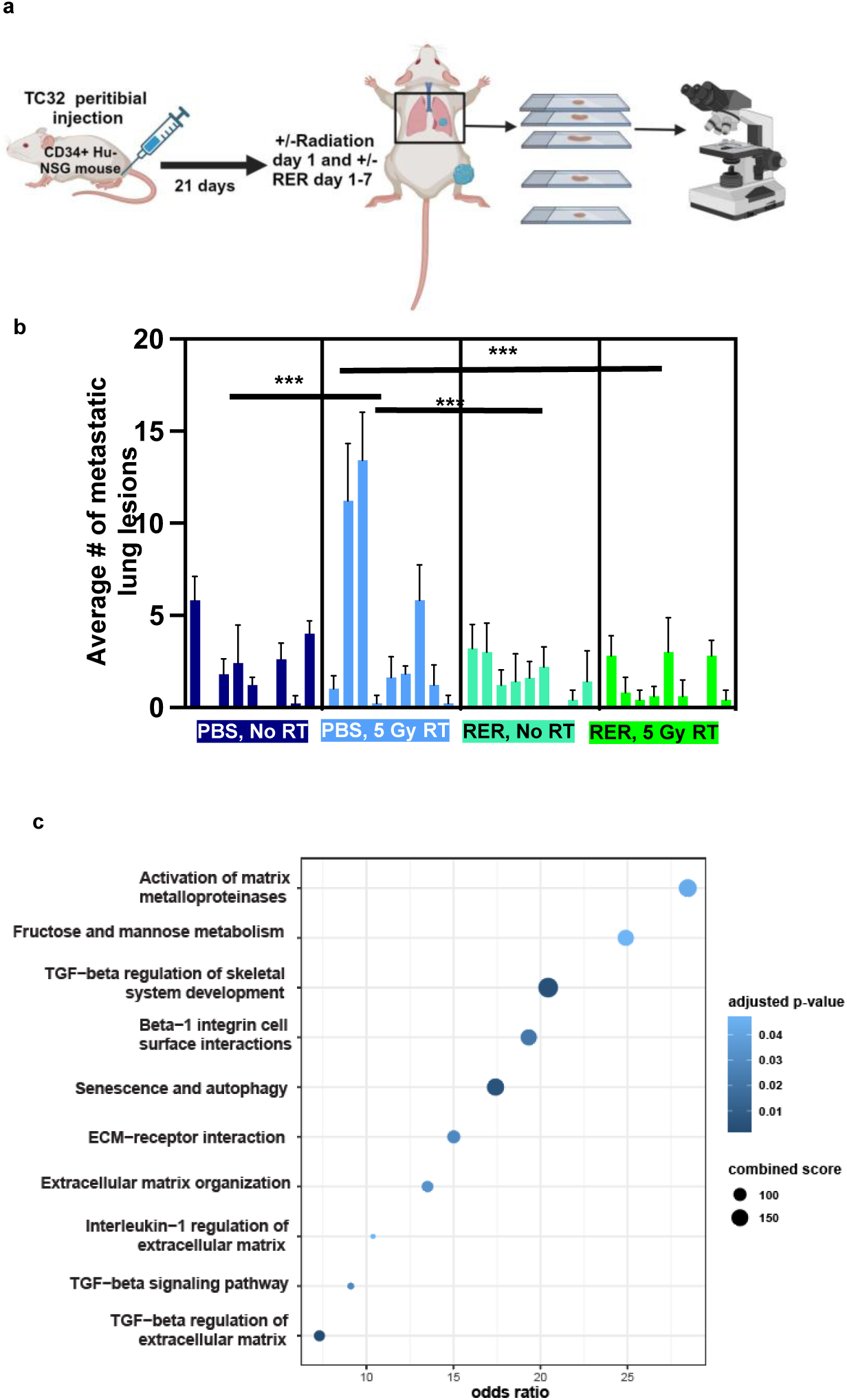
TGFβ inhibition during radiation suppresses the Ewing sarcoma metastatic phenotype. **a** Experimental overview of analysis for lung metastases. **b** Lungs were harvested from mice in the experiment from Fig 5. Lung tissue was processed for FFPE and underwent serial sectioning prior to review by independent pathologist. The number of metastatic foci per serial section was determined. Each bar represents an average number of metastatic lesions across serial section in individual mouse, error bars represent standard deviation. *** = p< 0.05. **c** Pathway analysis performed on the significantly differentially downregulated genes in Ewing tumors from mice treated with 5Gy RT and RER (TGFβ inhibitor) compared to treatment with 5Gy RT and PBS control utilizing EnrichR bioplanet 2019 library.

Lastly, to investigate potential mechanisms by which TGFβ inhibition reduces metastases following radiation in our model, we also analyzed the transcriptional profile of these tumors. Pathway analysis was performed on transcripts significantly downregulated in tumors that received radiation and TGFβ inhibition, compared to tumors receiving radiation alone, and reveals that 3 of the top 10 downregulated pathways involve TGFβ signaling as anticipated (Fig 6c). Additionally, significant downregulation of pathways involved in extracellular matrix organization and interactions (integrins) is noted. Taken together, these data demonstrate that TGFβ inhibition reduces tumor metastatic potential during radiation therapy and that pro-metastatic extracellular matrix genes may contribute to this effect.

## Discussion

Here, we demonstrate single-cell level expression patterns of TGFβ in human Ewing tumors, and then utilize an *in vivo*, immunocompetent humanized (hu-CD34+) mouse model to demonstrate that TGFβ inhibition enhances total immune cell infiltration and reduces lung metastases following radiation therapy in Ewing sarcoma. The major implications of these data include: 1) we have developed a new immunocompetent mouse model tool for the study of the Ewing sarcoma TME and the preclinical evaluation of TME modulating agents in Ewing sarcoma, 2) our data supports the potential use of TGFβ inhibition + radiation to promote immune cell infiltration into Ewing tumors, 3) our data indicates that TGFβ inhibition may reduce metastatic potential following radiation in Ewing sarcoma, and lastly, 4) this work serves to shift a historical dogma and highlight the important functional role of TGFβ in Ewing tumor biology. We will now discuss each of these points in greater detail.

First, the identification of appropriate pre-clinical models that can accurately predict the success of new therapies in early phase clinical trials remains a challenge (43, 44). Importantly, the role of the immune component of the tumor microenvironment in driving tumor response to therapy, and potential resistant mechanisms, is increasingly recognized as a crucial consideration in the design of new therapeutic strategies (7). There are no syngeneic or transgenic mouse models of Ewing sarcoma (17), so for this study we developed a humanized mouse model of Ewing sarcoma utilizing a hu-CD34+ model to be able to ask specific questions about how radiation therapy and TGFβ inhibition affect tumor immune response. Baseline Ewing tumor infiltration with immune cells in the hu-CD34+ model is low, similar to the low level of immune infiltrate noted in primary human Ewing tumors, including our own prior work (21). Our data clearly demonstrate transcriptional signature differences between identical Ewing tumors developed in hu-CD34+ versus NSG mice, thus highlighting the influence of immune cells on Ewing tumor biology. As humanized mouse models are very costly (∼4x the cost of experiments conducted in NSG mice), practical utilization of this model in the field should focus on studies addressing specific immunobiology questions or for the preclinical testing of agents that are thought to have potential tumor microenvironment or immunomodulatory effects. Incorporation of this model into pre-clinical testing of Ewing sarcoma may be important to appreciate immune-driven mechanisms of therapeutic resistance or efficacy.

Second, utilizing our hu-CD34+ orthotopic mouse model of Ewing sarcoma, we demonstrate that the combination of TGFβ inhibition with radiation therapy results in increased total immune cell (CD45+) infiltration into Ewing tumors. When considering practical applications of this finding, cellular therapies are an important consideration. A key barrier to the efficacy of cellular therapies for solid tumors, including Ewing sarcoma, is the limited infiltration of engineered cell therapies into the tumors (45–47). Our data suggest that priming tumors with TGFβ inhibition plus a single fraction of radiation prior to the delivery of cellular therapies may enhance their infiltration into the tumor. Testing this potential is a future direction of this work.

Third, little is currently understood regarding the mechanism(s) regulating Ewing sarcoma lung metastases (48). Our current findings demonstrate that more spontaneous lung metastases develop at baseline when comparing orthotopic Ewing tumors developed in hu-CD34+ mice versus NSG mice, suggesting there may an advantage to utilizing hu-CD34+ mice in studies specifically focused on Ewing lung metastases. We also demonstrate an increase in lung metastases in humanized mice following a single fraction of radiation therapy. While the exact mechanism of this effect is unknown, it is possible that radiation induces enhanced leakiness of the tumor vasculature (49), bleeding into the tumor that helps assist tumor cell escape, or immunosuppression. As we find TGFβ inhibition abrogates radiation-induced lung metastases, it is interesting to speculate on the potential clinical implications of this finding. Radiation therapy is utilized for the treatment of aggressive Ewing sarcoma (22, 23, 36), and these data provide support for additional preclinical combination studies of radiation and TGFβ inhibition for the treatment or prevention of Ewing lung metastases.

Lastly, historically, the impact of TGFβ in Ewing tumors has been assumed to be of limited consequence due to the repression of *TGFBR2 (*TGF-beta receptor 2) by the EWSR1::FLI1 oncoprotein (25). This thinking started to shift when the heterogeneity and plasticity of EWSR1::FLI1 expression in Ewing tumors was demonstrated (40, 41). Our current work demonstrates that TGFβ inhibition has a functional impact on Ewing tumor metastatic behavior following radiation in an *in vivo* immunocompetent context, highlighting the importance of considering the entire microenvironment when studying Ewing sarcoma biology. Future studies will aim to preclinically determine how to maximize the benefit of TGFβ inhibition during radiation therapy and potentially translate these findings to help prevent or treat lung metastases in patients with Ewing sarcoma.

## Acknowledgements

First and foremost, we would like to thank all the patients diagnosed with Ewing sarcoma and their families for their willingness to participate in research studies. Bulk RNAseq library generation and sequencing was performed by the University of Pittsburgh Health Sciences Sequencing Core (HSSC), Rangos Research Center, UPMC Children’s Hospital of Pittsburgh, Pittsburgh, Pennsylvania, United States of America. This work utilized the Hillman Cancer Center Biostatistics Facility, a shared resource at the University of Pittsburgh supported by the CCSG P30 CA047904. Single-cell RNAseq library preparation was conducted at The University of Pittsburgh Single Cell Core.

---

**Supplemental Figure 1.**
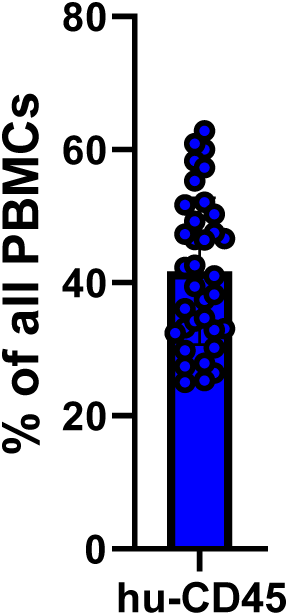
Human CD45+ cells present in our hu-CD34+ mouse model of Ewing sarcoma. Immunophenotyping for human CD45 was performed on peripheral blood mononuclear cells isolated from humanized mice. Error bars represent standard deviation.

**Supplemental Figure 2.**
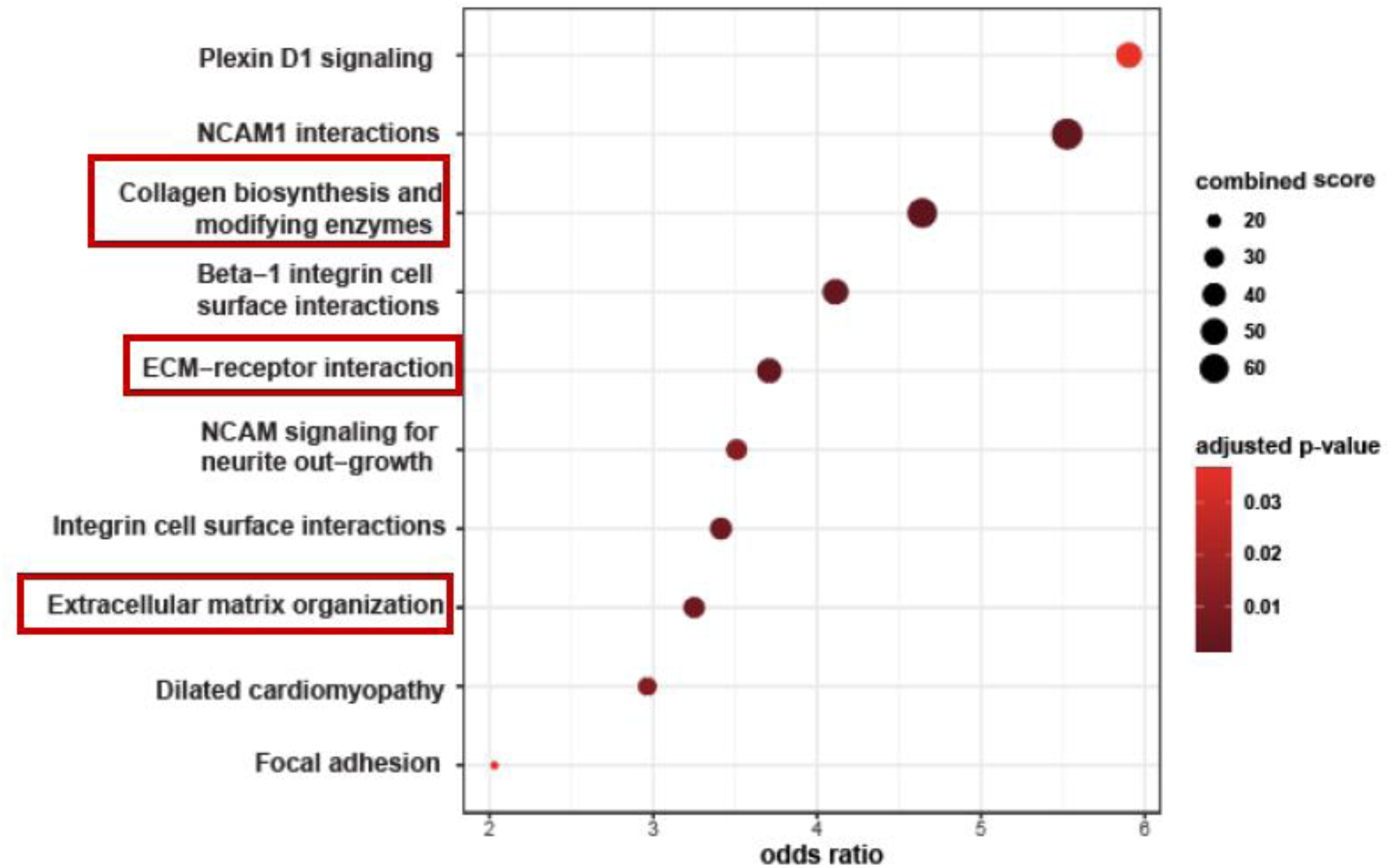
Ewing tumors developed in hu-CD34+ mice demonstrate upregulation of ECM pathways. Bulk RNA sequencing was performed on Ewing tumors developed in both hu-CD34+ and NSG mouse models. 2112 differentially expressed genes (p<0.05, FC >=1.5) were identified in tumors developed in hu-CD34+ model vs NSG model. Pathway analysis was performed on DEGs in the immunocompetent mouse model versus the immunodeficient model utilizing EnrichR bioplanet 2019 library.

**Supplemental Figure 3.**
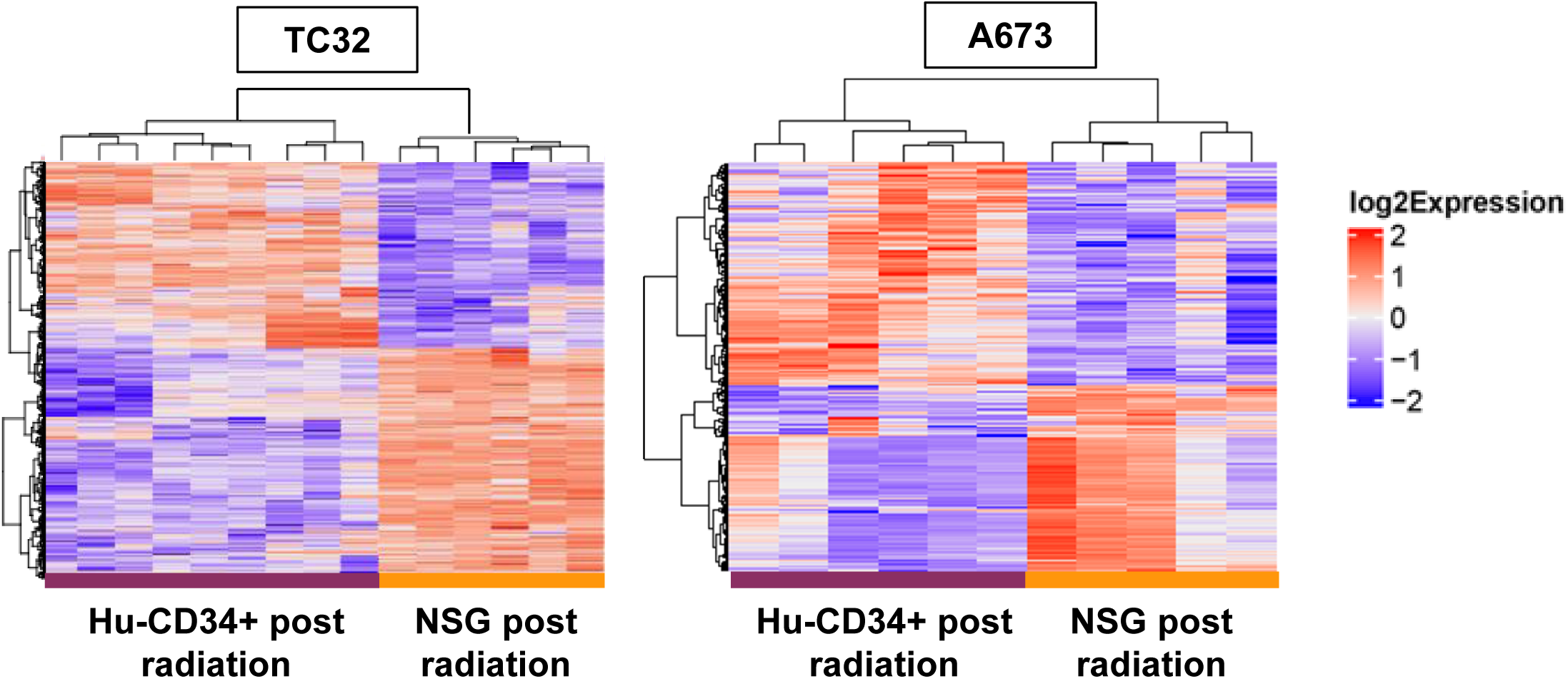
Distinct tumor transcriptional signatures following radiation therapy are noted when Ewing tumors are developed in hu-cD34+ versus NSG mouse models. Bulk RNA sequencing was performed on Ewing tumors (TC32 and A673) developed in immunocompetent (hu-CD34+) and immunodeficient (NSG) mice and treated with focal radiation therapy. Unsupervised clustering demonstrates that radiated tumors developed in hu- CD34+ mice or NSG mice cluster separately in both cell lines tested.

**Supplemental Figure 4.**
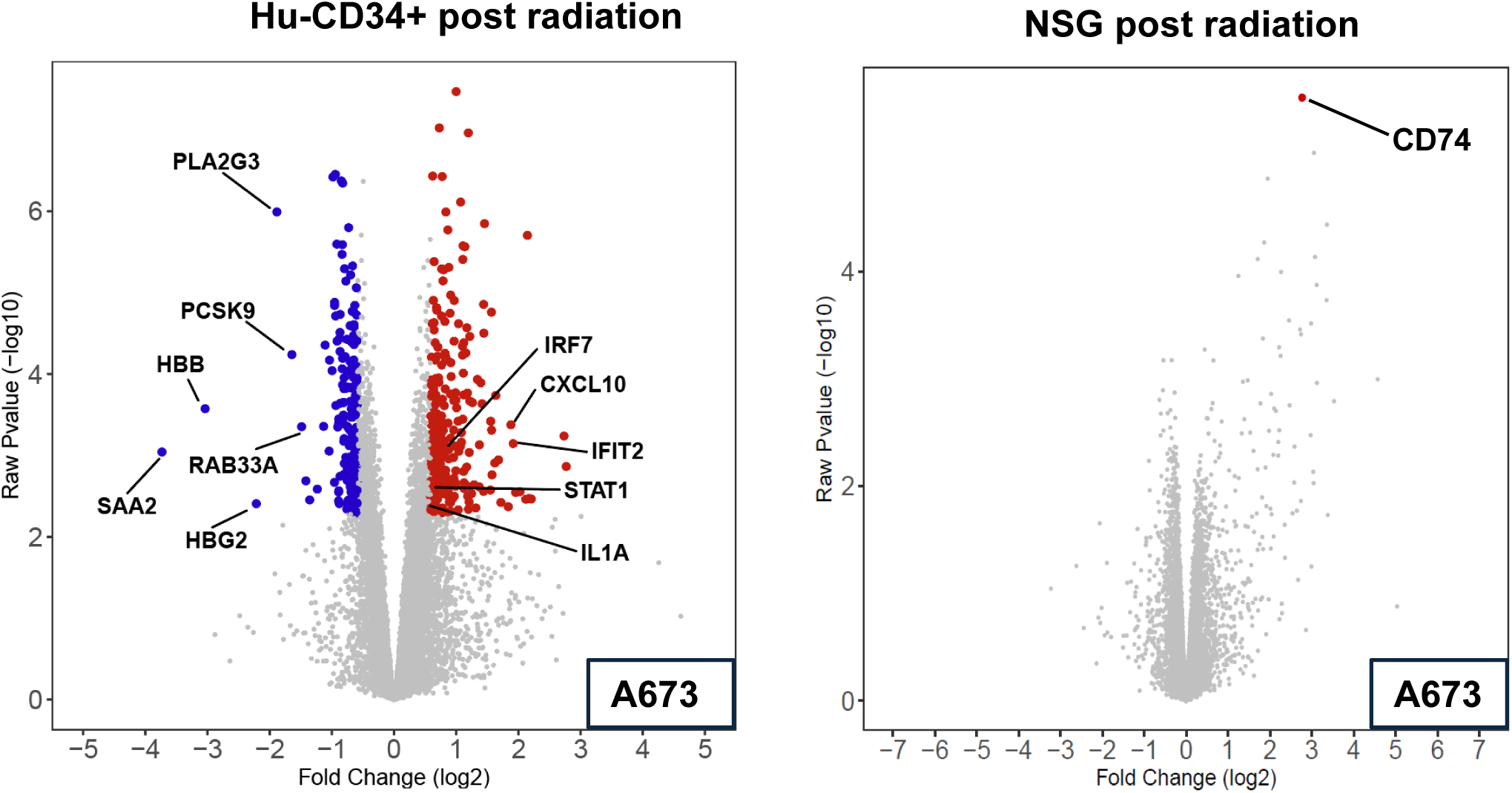
Transcriptional modulation induced following radiation treatment of A673 Ewing tumors developed in hu-CD34+ versus NSG mice. RNA was extracted from tumors post sacrifice and bulk RNA sequencing was performed. Volcano plots demonstrating differential gene expression following radiation treatment of A673 tumors derived in either hu-CD34+ or NSG mice are shown.

**Supplemental Figure 5.**
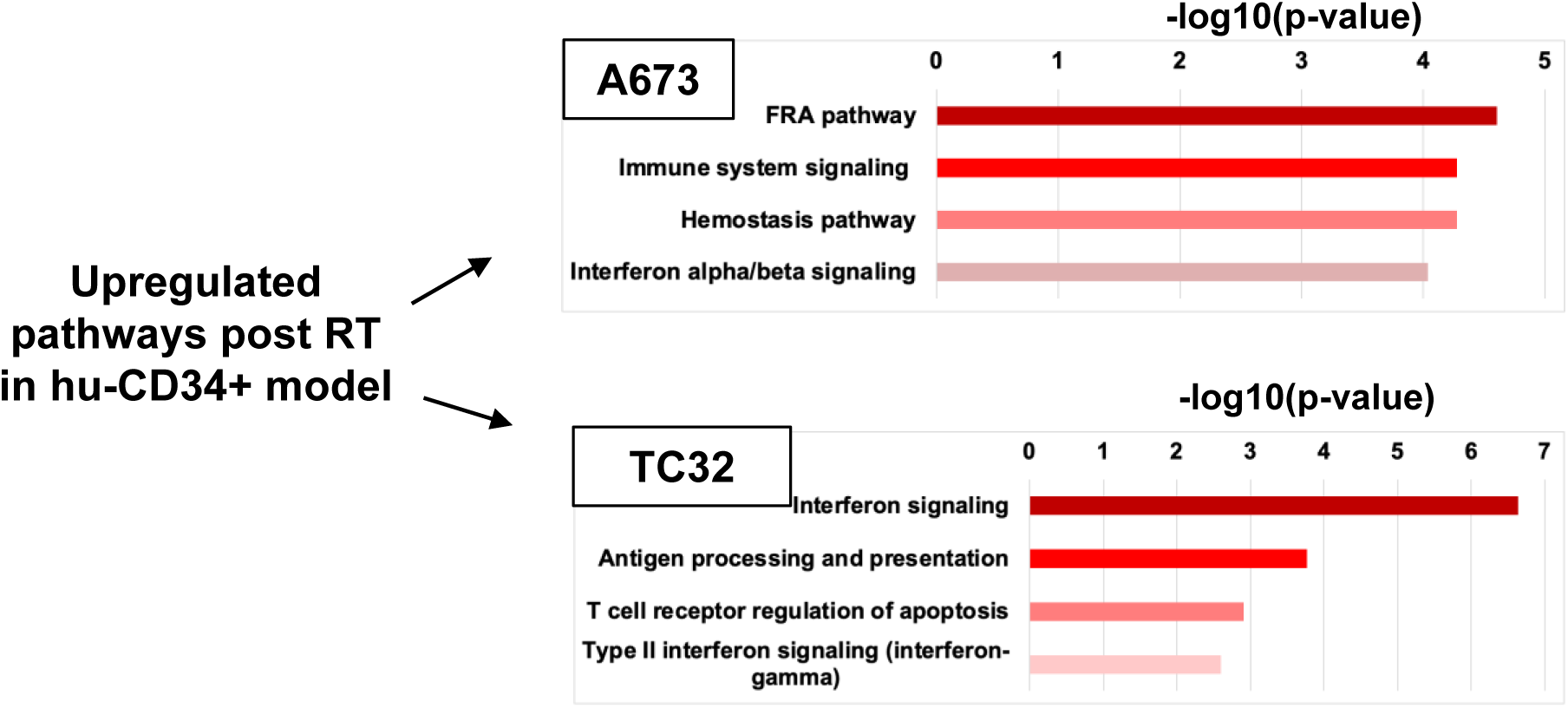
Radiation induces upregulation of inflammatory pathways in Ewing tumors developed in a hu-CD34+ mouse model. Pathway analysis was performed utilizing EnrichR Bioplanet 2019 on significantly differentially upregulated genes following radiation of tumors developed in hu-CD34+ mice demonstrating upregulation of pathways involved in inflammatory response.

**Supplemental Figure 6.**
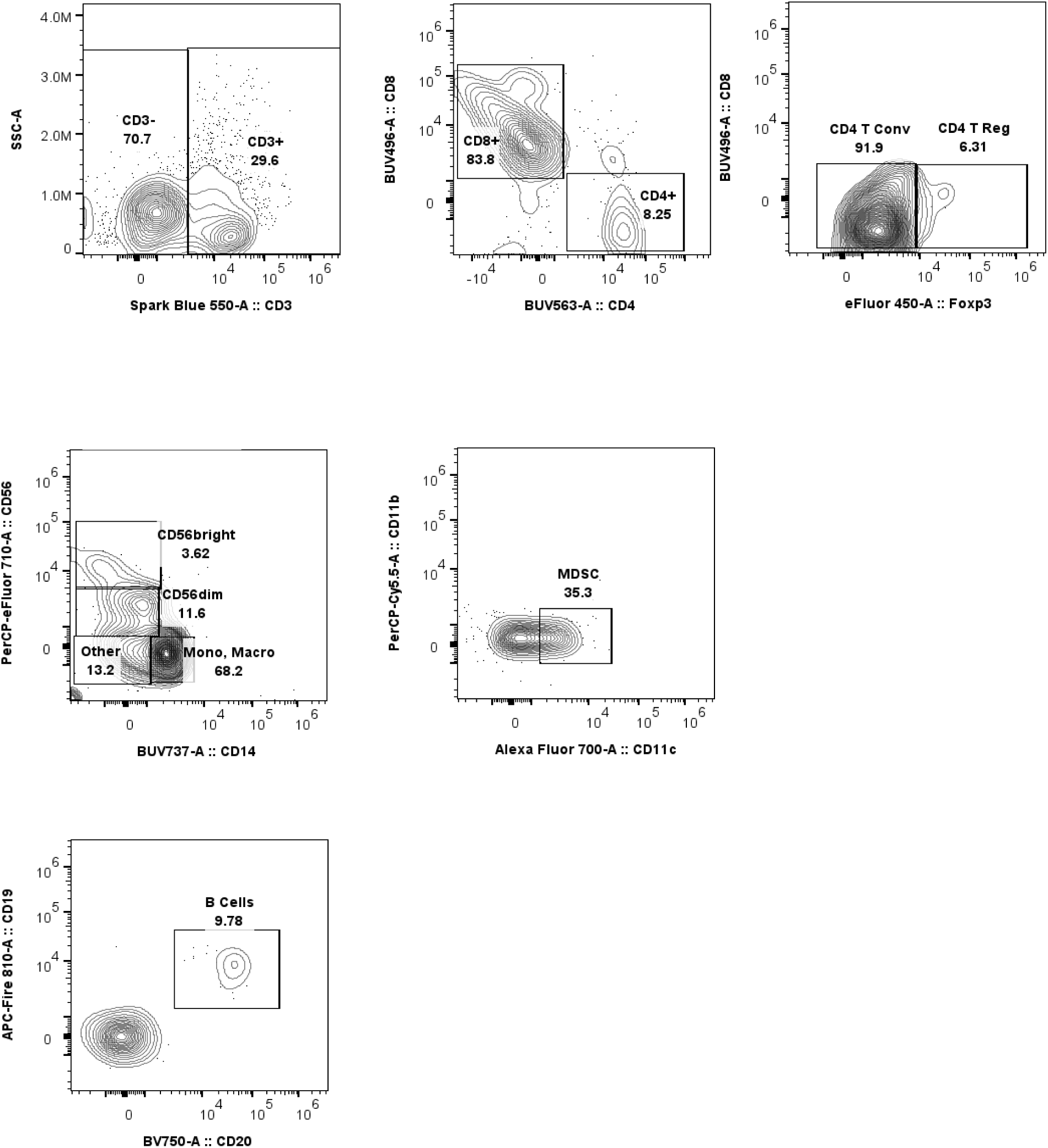
Flow cytometry gating strategy. Immunophenotyping was performed on single cell suspension of TC32 Ewing tumors developed in hu-CD34+ mice. Representative plots to demonstrate the gating strategy used to define immune cell sub-populations are shown.

**Supplemental Figure 7.**
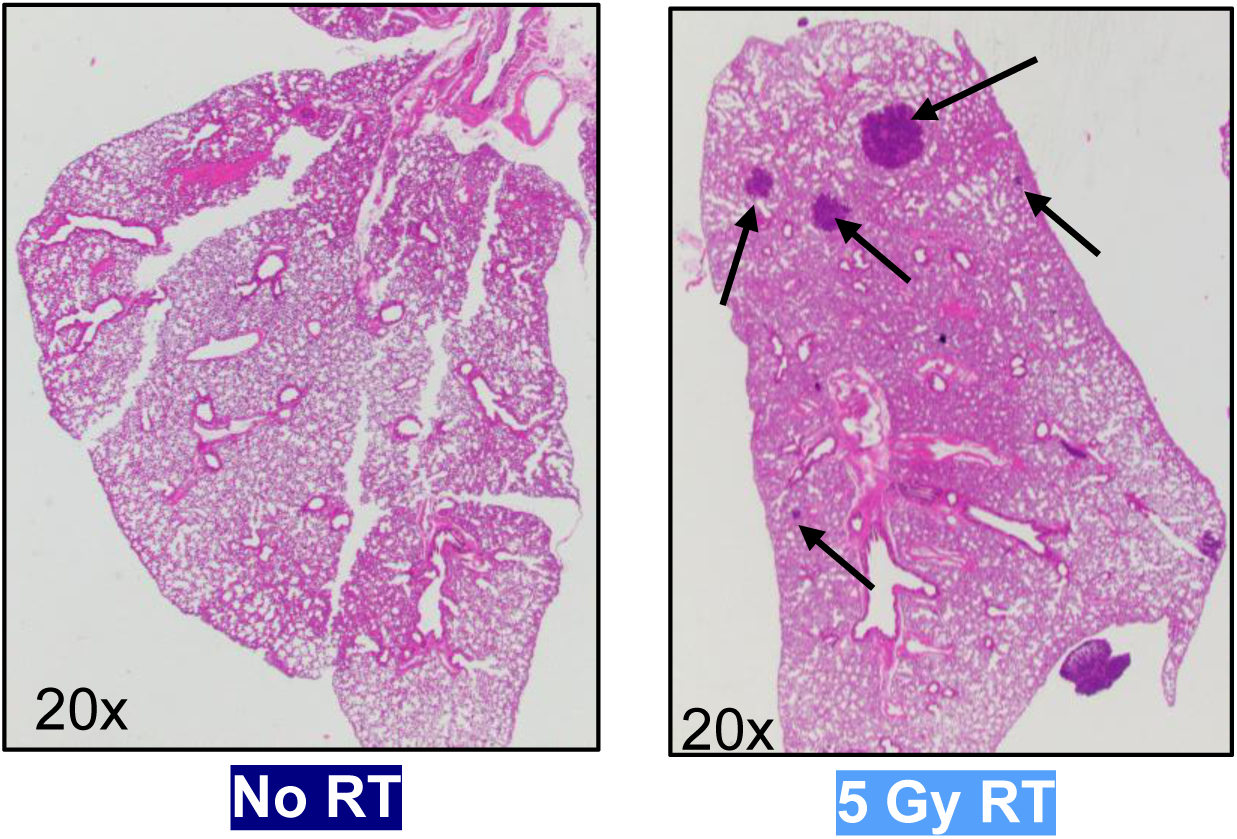
Representative H&E images of mouse lungs used for metastatic quantification. Representative images of lung tissue with no metastases (PBS, no radiation therapy), or with metastases noted (PBS, 5 Gy RT), see arrows. Corresponds to data shown in Fig 6b.

